# Systemic delivery of cationic liposome-mediated siRNA EGFR enhances therapeutic efficacy in a human colorectal cancer model

**DOI:** 10.64898/2026.03.29.715100

**Authors:** Damian Kaniowski, Anna Boguszewska-Czubara, Katarzyna Ebenryter-Olbińska, Katarzyna Kulik, Justyna Suwara, Artur Wnorowski, Jakub Wójcik, Barbara Budzyńska, Agnieszka Michalak, Algirdas Ziogas, Barbara Nawrot, Olga Swiech

**Affiliations:** Biotechna S.A., Dobrzanskiego 3, 20-262 Lublin, Poland; Centre of Molecular and Macromolecular Studies, Polish Academy of Sciences (CMMS PAS), Sienkiewicza 112, 90-363 Lodz, Poland; Department of Medical Chemistry, Medical University of Lublin, Chodzki 4A, 20-093 Lublin, Poland; Department of Biopharmacy, Medical University of Lublin, 20-093 Lublin, Poland; Faculty of Chemistry, University of Warsaw, Pasteura 1, 02093 Warsaw, Poland; Independent Laboratory of Behavioral Studies, Medical University of Lublin, 20-093 Lublin, Poland; Faculty of Fundamental Sciences, Vilnius Tech University, Vilnius, Lithuania

**Keywords:** cationic lipids, control release, therapeutic nucleic acids, siRNA, EGFR, targeted delivery, colorectal cancer

## Abstract

The clinical translation of RNA interference (RNAi) therapeutics remains limited by inefficient delivery and cancer-target accumulation. Here, we report the development of a new cationic liposome (CLP) nanocarrier engineered for delivery and controlled-release of small interfering RNA (siRNA) targeting the epidermal growth factor receptor (EGFR) in human colorectal cancer. CLPs were synthesized from ethylphosphocholine-based lipids and PEGylated components, with folic acid (FA) tissue-specific ligand and fluorophore labelling. These nanocarriers exhibited robust physicochemical stability across a broad pH and temperature range, efficient siRNA complexation, and nuclease-protection of siRNA. Functional studies revealed that CLP-siEGFR achieved effective cytosolic siRNA cargo release and EGFR silencing in vitro, proving to be more effective than conventional lipid-based transfection systems. In human xenograft models, intravenously administered CLP-siEGFR showed enhanced tumor localization, prolonged siRNA retention, and significant tumor growth suppression, accompanied by marked downregulation of EGFR. Importantly, systemic dosing was well-tolerated, with no evidence of hepatotoxicity, nephrotoxicity, or hematological abnormalities. These results position CLP nanocarriers as an effective platform for targeted RNAi therapeutics, offering translational potential for precision oncology applications.

## 1. Introduction

Colorectal cancer (CRC) is the third most diagnosed malignancy worldwide, with projections estimating an alarming rise to 3.2 million new cases and 1.6 million fatalities by 2040.^1^ Notably, up to 25% of patients with CRC are diagnosed with stage IV disease, while up to 50% of those initially presenting with early-stage CRC progress to metastatic disease over time. The prognosis for stage IV CRC remains dismal, with a 5-year survival rate of only 12.5% in the United States and 15.4% in Europe.^2,3^ These statistics highlight a critical unmet clinical need for therapeutic strategies that combine high drug delivery efficacy with a favorable safety profile to effectively address the growing burden of colorectal cancer.^4^

Epidermal growth factor receptor (EGFR) aberrant in colorectal cancer is predominantly driven by protein overexpression, which is observed in approximately 35% to 50% of patients. Numerous studies have established a statistically significant correlation between EGFR overexpression and adverse prognostic outcomes.^5^ The EGFR signalling pathway is integral to cancer progression, leading to the development and clinical introduction of several targeted therapies, including cetuximab and panitumumab.^6^ These monoclonal antibodies competitively bind to the extracellular domain of EGFR, preventing receptor tyrosine kinase phosphorylation and activation.^7^ This mechanism suppresses cell growth, induces apoptosis, and promotes cell cycle arrest. Moreover, EGFR activation can promote tumor growth by promoting VEGF upregulation through a hypoxia-independent mechanism.^8^ Additionally, cetuximab can stimulate antibody-dependent cellular cytotoxicity, contributing to immunogenic cell death and enhancing the immune-mediated effects of therapy.^8,9^ These targeted therapies have demonstrated substantial efficacy in selected subsets of CRC patients, significantly slowing disease progression.^10,11^ Comprehensive preclinical research on the mechanisms of resistance to anti-EGFR monoclonal antibodies has underscored the critical need for the development of new and EGFR-targeted therapies.^12^

Despite advances in the development and clinical application of 1^st^, 2^nd^ and 3^rd^ generation EGFR tyrosine kinase inhibitors (TKIs), drug resistance remains a significant challenge. Clinical trials have highlighted issues such as inadequate tumor accumulation, limited penetration into solid tumor masses, and impediments posed by the tumor microenvironment.^13^

To overcome current limitations in clinical translation, therapeutic nucleic acids represent a powerful strategy for targeted therapy. Small interfering RNAs (siRNAs) are short double-stranded RNA molecules, typically 19-25 base pairs in length, capable of integrating into the RNA-induced silencing complex (RISC) to induce sequence-specific silencing of target mRNA.^14^ To date, several siRNA-based therapeutics have received FDA approval for clinical use, targeting gene expression in rare metabolic and cardiovascular disorders. These include Patisiran (Onpattro) for hereditary transthyretin-mediated amyloidosis (hATTR); Givosiran (Givlaari) for acute hepatic porphyria (AHP); Lumasiran (Oxlumo) and Nedosiran (Rivfloza) for primary hyperoxaluria type 1 (PH1); Inclisiran (Leqvio) for lowering LDL cholesterol in patients with atherosclerotic cardiovascular disease (ASCVD); and Vutrisiran (Amvuttra), also for hATTR and Fitusiran (Qfitlia) for hemophilia.^15,16^ These therapies utilize either lipid nanoparticle or N-acetylgalactosamine ligand (GalNAc)-conjugate delivery systems to achieve targeted, durable gene silencing via RNAi.^17^ Despite siRNA therapeutic potential, several significant challenges persist for the in vivo application of siRNA, including off-target effects, inefficient delivery with poor cellular uptake, and activation of immune responses, all of which hinder their clinical feasibility.^18^ These obstacles highlight the need for more efficient and targeted delivery systems to enhance the effectiveness of siRNA-based therapies. Various delivery technologies and strategies, including lipophilic conjugates, cationic polymer- or micelle-based carriers, protein-antibody conjugates, and modifications to the size and structure of the siRNA molecule itself, are under development. These approaches have demonstrated robust efficacy in different animal models, with some platforms advancing toward clinical translation.^14,19,20,21^ Lipid nanoparticle delivery was the first successful strategy for gene silencing in human and enabled the approval of the first siRNA-based therapeutic.^22,23^

Despite clinical progress, the use of RNAi nanomedicines in the treatment of solid tumors remains limited by low tumor accumulation and inefficient intracellular transport^24^. Quantitative meta-analyses indicate that on average only about 0.7% of the injected dose (ID) of nanoparticles accumulates in solid tumors, with a median of 0.6% ID, highlighting limitations related to heterogeneous vascular permeability and clearance by the mononuclear phagocyte system.^25,26^ A second, well-recognized bottleneck is endosomal entrapment. Image-based single-cell tracking has shown that only about 1-2% of internalized siRNA molecules escape from endosomes into the cytosol, with the vast majority remaining trapped and ultimately degraded in endo-lysosomal compartments.^27^

Polyethylene glycol (PEG)-lipids are widely used to minimize aggregation and opsonization; however, they introduce the well-known ‘PEG dilemma’. While PEG prolongs systemic circulation, excessive surface shielding can impede cellular uptake and endosomal escape. Furthermore, repeated administration may lead to the accelerated blood clearance (ABC) phenomenon.^28,29^ Beyond PEG density, the terminal-group chemistry of PEG is emerging as a tunable parameter that can modulate interfacial charge, protein corona composition, and tumor retention. Evidence from reviews and studies on targeted liposomes indicates that functional PEG end-groups (e.g., folate or thiol-reactive moieties) can enhance tumor cell interaction compared to inert methoxy-PEG.^30,31^ Building upon this rationale, we employed our proprietary PEG-functionalized cationic liposome (CLP) platform (WIPO WO2023022615). Incorporation of PEG-NH₂ lipids combined steric stabilization with enhanced electrostatic association of siRNA, preserving a positive surface charge without inducing aggregation and enabling release modulation through surface chemistry. This system enabled selective accumulation of siRNA inhibitor within colorectal tumor tissue, resulting in efficient *EGFR* gene silencing and significant tumor growth inhibition. The CLP formulation was well tolerated in vivo, with no evidence of systemic toxicity. These findings support the new CLP platform as a promising candidate for further preclinical translation in RNAi-based precision oncology and clinical translation.

## 2. Results

### 2.1. Design and optimization of cationic liposomes for RNAi-based applications

Cationic liposomes (CLP) developed in this study were formulated using saturated cationic ethylphosphocholines (EPC) in combination with saturated neutral lipids (DPPC and DMPC) and polyethylene glycol (PEG)-modified lipids bearing a terminal primary amine group. EPCs are cationic derivatives of phosphatidylcholine, where the phosphate oxygen’s negative charge is neutralized by the addition of an ethyl group. This modification results in a compound that is chemically stable, biodegradable, and composed entirely of biological metabolites linked by ester bonds. The CLP were synthesized via the conventional thin-film hydration method, followed by extrusion to ensure uniform size distribution (Figure 1A). This approach enabled the preparation of liposomes with a diverse range of lipid compositions, varying in charge, fatty acid chain length, and surface modifications with either a targeting folic acid-ligand (FA) or a Cyanine 7 dye (Cy7). For this study, cationic liposomes CLP and folic acid-modified cationic liposomes (CLP-FA) were selected as representative formulations, incorporating EPC lipids and PEG-modified lipids with terminal amine functionality (Figure 1B). These formulations were developed and described in our patent applications (PCT.PL2022.000046; WO 2023/022615 A1). The synthesized CLPs, CLPs-FA, and their Cy7-labeled counterparts exhibited small hydrodynamic sizes and low polydispersity indices (PDI), indicative of a monodisperse population. Notably, the incorporation of DSPE-PEG(2000)-folate lipid and 1,2-distearoyl-sn-glycero-3-phosphoethanolamine conjugated with Cyanine 7 (18:0 Cy7-PE) lipid did not induce significant alterations in either size or PDI.

**Figure 1.**
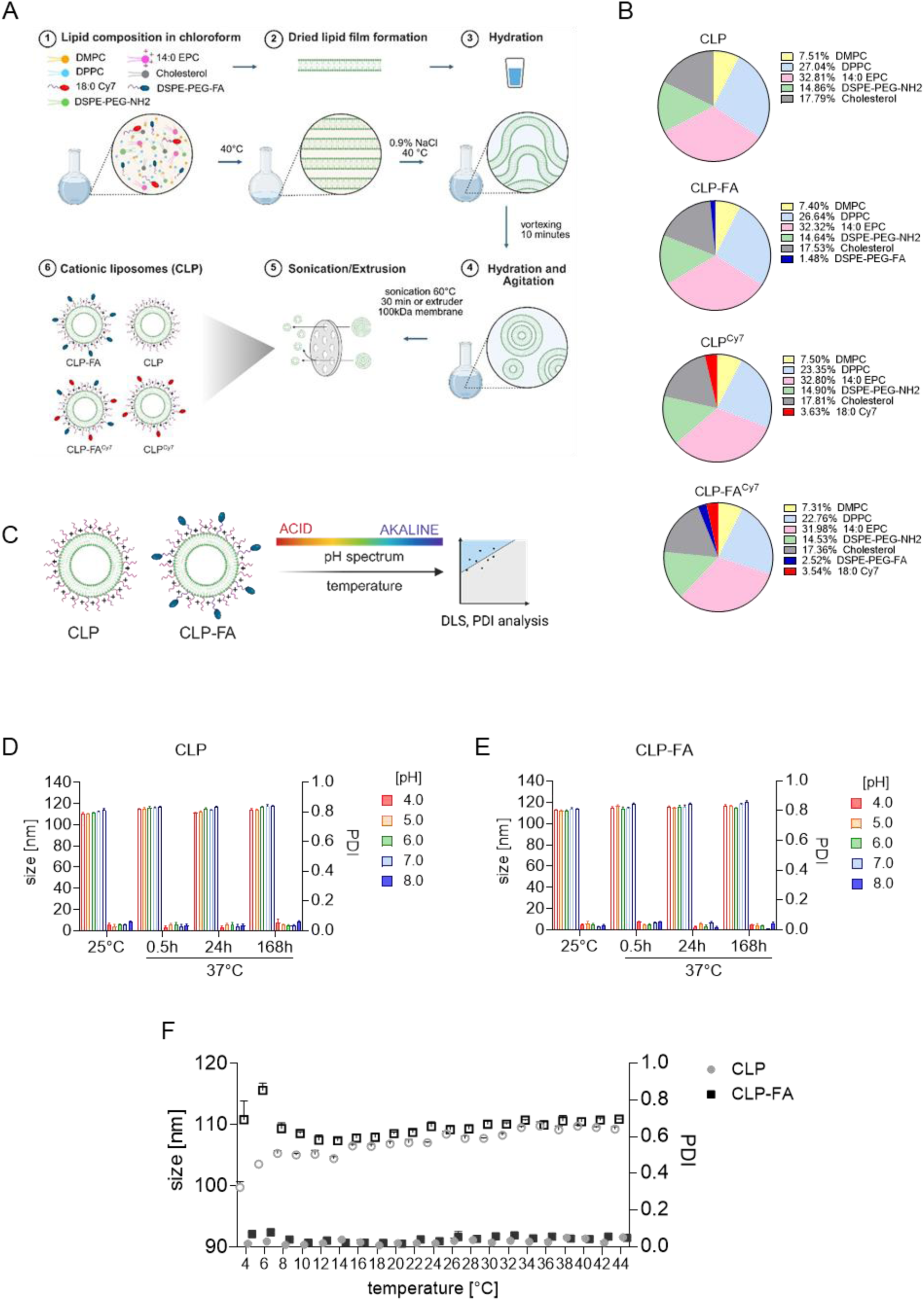
Engineering the composition of cationic liposomes for the delivery of siRNA. Schematic representation of the synthetic procedure for CLPs liposomes (**A**); a pie chart showing the percentage composition of individual lipids (**B**); stability studies (**C**); changes in hydrodynamic radius (size, empty bars) and PDI (filled bars) of CLP and CLP-FA in electrolytes with varying pH levels (pH 4.0 to 8.0) at 25°C or after incubation at 37°C for up to 168h (7 days) (**D**, **E**); the temperature profile of liposomes across a range of 4°C to 44°C. The size (upper empty rectangles and circles) and PDI (lower filled rectangles and circles) (**F**).

To assess the stability of the liposomal formulations, temperature profile analysis and physicochemical evaluations across a pH range of 4.0 to 8.0 were conducted (Figure 1C). Both CLP CLP-FA liposomes demonstrated stability across a wide range of pH values and temperatures (Figure 1D, 1E). Stability assessments revealed that the hydrodynamic sizes of CLP and CLP-FA remained constant, with PDI values consistently below 0.1, in PBS buffers ranging from pH 4.0 to 8.0 (Figure 1D, 1E, respectively). Furthermore, these parameters remained unchanged at room temperature (25°C) and after one week of incubation at 37°C, underscoring the structural integrity of the liposomes under physiological conditions relevant for intravenous administration. The thermal stability of the liposomes was further examined across a temperature range of 4°C to 44°C (Figure 1F). Throughout this range, PDI values remained below 0.1, and no significant size variations were observed, suggesting the absence of lipid bilayer phase transitions in the selected lipid composition. These findings confirm that CLP and CLP-FA maintain their structural integrity under physiologically relevant conditions. Long-term stability studies have shown that both CLP and CLP-FA remain stable under long-term storage conditions (Figure S1 A-C).

### 2.2. RNAi-based therapeutic targeting of EGFR in colorectal cancer

Kaplan-Meier survival analysis demonstrated that colon cancer patients with high EGFR expression exhibited significantly poorer prognosis compared to those with low EGFR expression, highlighting the clinical relevance of targeting this receptor as a potential therapeutic strategy (Figure 2A). The extracellular domain of EGFR is currently targeted by monoclonal antibodies (e.g., cetuximab) in clinical settings; however, these therapies have shown limited efficacy in certain patient cohorts.^32^ To address this limitation, we designed a small interfering RNA (siRNA) specifically targeting the mRNA of EGFR within the sequence encoding domain III (L2) of EGFR, which is critical for ligand (EGF) binding (Figure 2B). To evaluate efficacy, siRNA EGFR (siEGFR) inhibitors were chemically unmodified or modified with 2′-*O*-methyl (siEGFR^2′-OMe^) at the ribose 2′-position, and with or without additional terminal stabilization using phosphorothioate-modified (siEGFR^PS^) internucleotide linkages (Figure 2C). All sequences and chemical modification patterns of the oligonucleotides used are presented in Figure 2D. siRNA compounds were purified from reaction mixtures using standard HPLC procedures and characterized by electrospray ionization quadrupole time-of-flight mass spectrometry (ESI-Q-TOF MS) (Figure S2).

**Figure 2.**
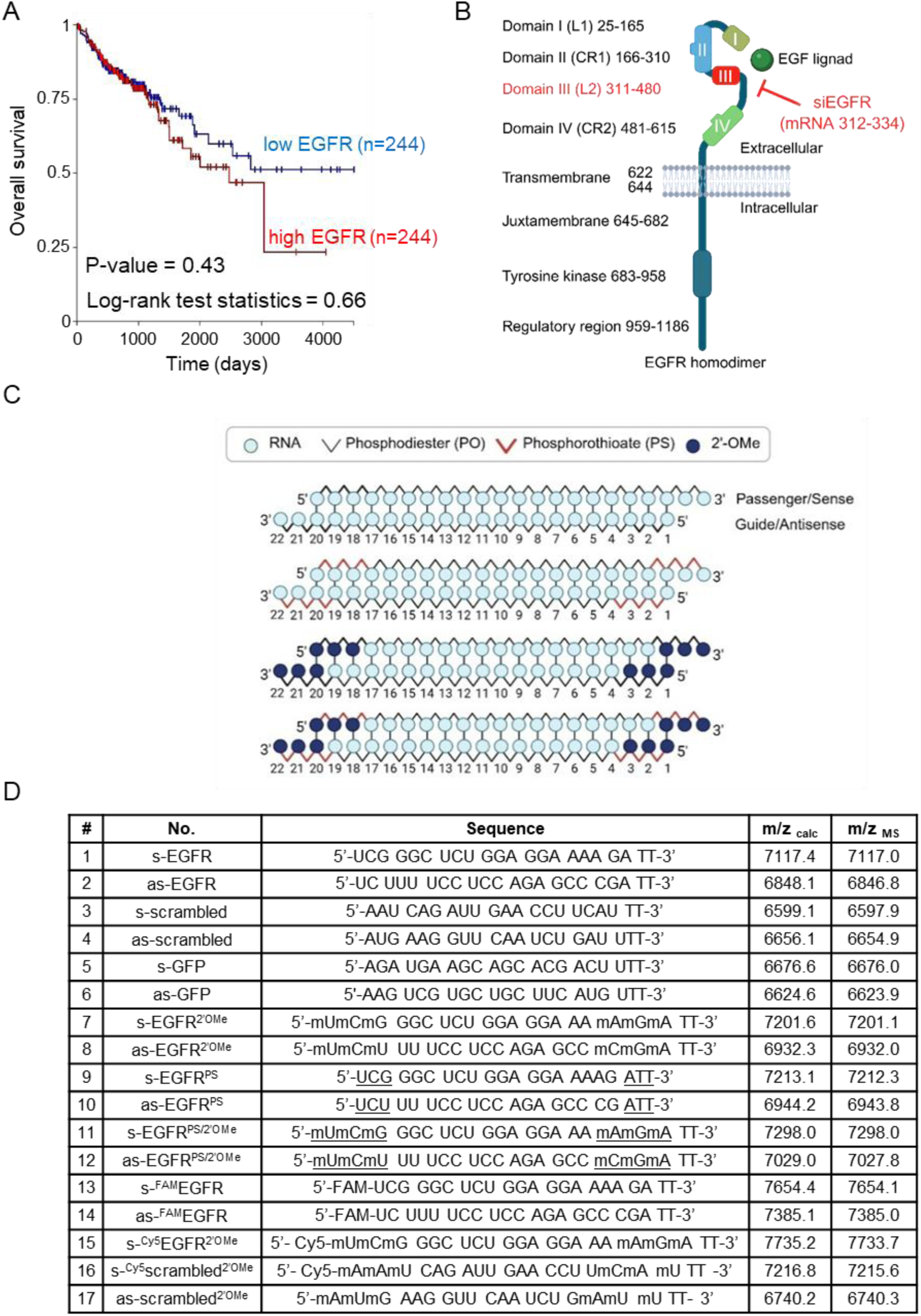
Design and characterization of siRNA targeting EGFR ligand-binding domain. Kaplan-Meier curves for overall survival of human colon cancer (GDC TCGA COAD) according to human EGFR categorized by gene expression RNAseq using UCSC Xena database. Survival time is presented in days; p values are related to Log-rank test results (**A**), homodimer structure of human EGFR based on UniProt: P00533 with EGF binding domain III. Screened siEGFR targets mRNA within the coding region of this domain (nucleotides mRNA 321-334; red) (**B**); Chemical modification patterns abbreviations: 2’-O-methyl (2’-OMe) - navy blue, ‘_’-phosphorothioate bonds (PS) - red, phosphodiester bonds (PO) - black, ribonucleic acid (RNA) – blue, 6-carboxyfluorescein dye (FAM) and Cyanine5 dye (Cy5) (**C**) and table of all RNA sense (s) and antisense (as) sequences (**D**).

### 2.3. Cationic liposome-mediated siRNA delivery overcomes the limitations of endosomal escape

CLP have long been recognized as highly effective gene delivery vehicles, demonstrating efficient biodegradability, biocompatibility, and high nucleic acid encapsulation rate.^33,24^ Zeta potential analysis provided further mechanistic insights into siRNA-lipid interactions (Figure 3A). The surface charge of CLP nanoparticles was monitored following the addition of either unmodified siEGFR (more hydrophilic) or modified siEGFR^2’-OMe^ (more hydrophobic, Figure 3B). Pristine CLPs exhibited a zeta potential of +66.7 mV. Upon complexation with siRNA at increasing siRNA:CLP mass ratios (1:100, 1:50, 1:20, 1:10), a progressive reduction in zeta potential was observed for both siRNA variants. Notably, the decrease was consistently less pronounced for 2′-*O*-methyl modified siRNA, ranging from ∼8% at a 1:100 ratio (+58.2 mV vs. +63.9 mV for unmodified and modified siRNA, respectively) to ∼25% at a 1:10 ratio (+28.9 mV vs. +39.0 mV, respectively) in Figure 2C. PDI analysis confirmed the colloidal stability of the formulations. Both unmodified and modified siRNA induced only minor increases in PDI (0.07 to 0.19), excluding aggregation and supporting the interpretation that siRNA is incorporated within the CLP interior and outer PEG layer without extensive interparticle bridging (Figure 3C). Comparable zeta potential trends were observed for folate-modified CLPs (Figure S1 A-C), indicating that siRNA binding follows a conserved mechanism independent of surface functionalization.

**Figure 3.**
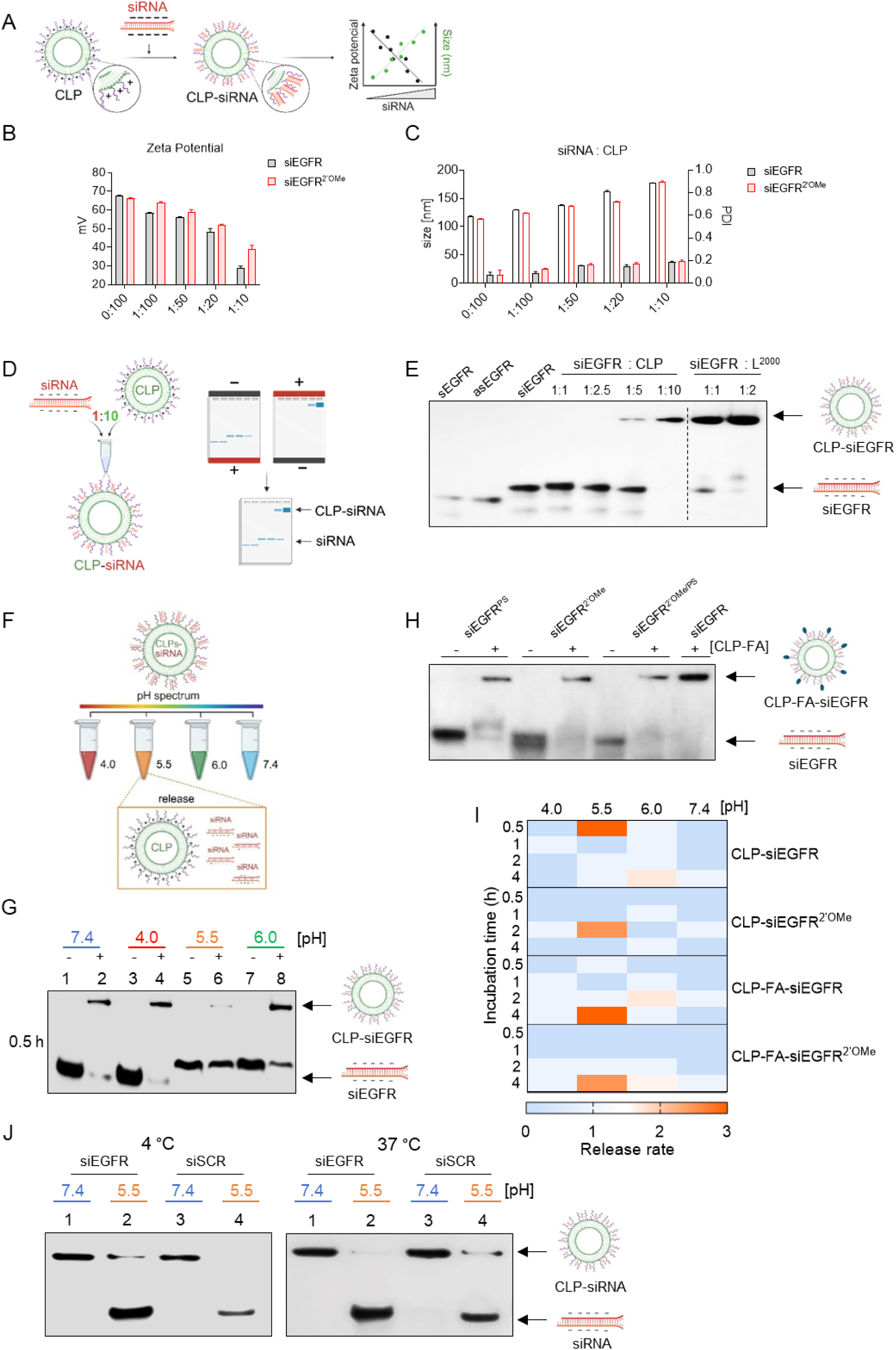
siRNA-loaded cationic liposomes facilitate efficient endosomal escape-associated intracellular delivery. Scheme of the study on siRNA binding to CLP cationic liposomes and readouts (**A**); Effect of siRNA loading at different ratios on the zeta potential of CLP (**B**); hydrodynamic radius (size, empty bars) and PDI (filled bars) after siRNAs addition on the CLP (**C**); Schematic showing binding of 5′-[³²P]-labeled siRNA to cationic CLPs, analyzed by PAGE in both directions (**D**); 5′-[^32^P]-labeled siRNA binding to CLP or Lipofectamine 2000 (L^2000^) at various mass ratios (w:w), sEGFR and asEGFR are single-stranded siRNA components (**E**); Schematic of pH-dependent siRNA release from CLP nanocarriers (lanes 2, 4, 6 and 8) and control siRNA (lanes 1, 3, 5, 7) across physiological and endosomal conditions (pH 7.4, blue; 6.0, green; 5.5, orange; 4.0, red) (**F, G**); Binding efficiency of chemically modified (2′-OMe, PS, and 2′-OMe/PS) and subsequently 5′-[^32^P]-labeled siRNAs to CLP-FA after 30 min incubation (**H**); Heat Map of siRNA release from CLP and CLP-FA at pH 4.0-7.4 over time were calculated by densitometry (**I**); CLP-siRNA complex stability was evaluated after 7 days of incubation at 4 °C and 37 °C (lanes 1-4) at pH 7.4 and under controlled-release conditions at pH 5.5 (lanes 2 and 4) (**J**); Analysis were performed using 15% PAGE under non-denaturing conditions.

Quantitative binding assays using 5′-[^32^P]-labeled siRNA and CLPs were performed to determine the stoichiometry of siRNA-CLP interactions (Figure 3D). Subsequently, the binding of negatively charged [^32^P]-siRNA to positively charge CLP was evaluated using PAGE analysis in both directions-from the negative to the positive terminal to visualize the single RNA or siRNA mobility and in the reverse direction to observe the complex (Figure 3D). The results indicated that the [^32^P]-siRNA exhibited optimal binding to CLP at a charge mass ratio of 1:10 (Figure 3E). The commercial transfection reagent Lipofectamine 2000 was used as a positive control (Figure 3E).

Endosomal escape refers to the crucial process by which nucleic acid-loaded nanoparticles are released from endosomes-acidic compartments with a pH ranging from 5.5 to 6.5-into the cytosol, which maintains a neutral pH of approximately 7.4.^34^ siRNA release from the CLP-siRNA formulation was assessed using acidified solutions that closely mimic the conditions of endosomal-lysosomal pathway (Figure 3F). Our results demonstrated that the CLP platform efficiently releases siRNA within the endosomal pH range of 5.5 to 6.5 after 30 min of incubation (Figure 3G). Moreover, the CLP-siRNA complex remains stable at pH 4.0 and 7.4. Next, we evaluated the binding efficiency of chemically modified siRNAs (2′-OMe, PS, and 2′-OMe/PS) to the cationic folic acid-liposome (CLP-FA) platform and found comparable binding affinity to that of unmodified siRNA EGFR (Figure 3H). Heat map analysis revealed that both modified and unmodified siRNAs are released from CLP or CLP-FA within the pH range of 5.5 to 6.0, but not at other pH levels (Figure 3I). Interestingly, the release duration of siRNA was extended to 4-hours in the CLP-FA platform compared to unmodified CLP (Figure 3I). To confirm that CLP does not degrade unmodified siRNA across sequences (EGFR and scrambled), stability analyses were conducted after 7 days of incubation at 4°C and 37°C using PAGE under release conditions (pH 5.5). As shown in Figure 3J, the CLP platform supports siRNA stability over one week across temperatures.

### 2.4. CLP-mediated EGFR siRNA delivery effectively reduces colorectal cancer growth

To evaluate uptake of the siRNA-CLP formulation by human colorectal cancer Caco-2 cells, we generated a fluorescently labeled EGFR-targeting siRNA (CLP-FA-siEGFR^FAM^). Prior to experiments, we assessed binding of siEGFR^FAM^ to CLPs and confirmed stable complex formation for up to 24 h (Figure S3B). Fluorescence microscopy confirmed the efficient and selective uptake of CLP-FA-siEGFR^FAM^ by Caco-2 cells in vitro after 30 minutes of incubation, in contrast to normal mouse fibroblasts (MEF-WT) (Figure 4B). The control experiment included transfection of siEGFR^FAM^ using Lipofectamine demonstrated comparable uptake of labeled siRNA (Figure S3A). Comprehensive evaluation of Cy5-labeled CLP-siEGFR^2′-OMe^ demonstrated concentration-dependent uptake in Caco-2 cells (Figure 4C) and confirmed complete carrier loading (Figure S3B). Both CLP and CLP-FA platforms without siRNA cargo showed no cytotoxicity after 48 h of incubation in Caco-2 cells, normal human colon epithelial cells (CCD-841CoN) (Figure 4D) and macrophages (Figure S3C), relative to the positive control. Next, to evaluate the EGFR gene-silencing activity of CLP-FA-siEGFR, we used a dual fluorescence assay (DFA) previously developed in our laboratory.^35,36,37^ We confirmed that the CLP platform efficiently inhibited the EGFR target by approximately 65% using modified siEGFR (2’-OMe and PS) and unmodified siRNA in Caco-2 cells, compared to the CLP-scrambled negative control (CLP-SCR), as shown in Figure 4F. In contrast, CLP-FA-siEGFR showed slightly reduced potency but still suppressed EGFR expression by 40% in Caco-2 cells relative to CLP-SCR (Figure 4G). In both platforms, GFP-targeting siRNA served as a positive control and effectively reduced EGFR-GFP fluorescence in Caco-2 and A431 cells (Figure S3 D-E), while naked panel of siRNAs were transfected using Lipofectamine (Figure S3 F-G). Furthermore, flow cytometry demonstrated that CLP-siEGFR^2′-OMe^ markedly reduced EGFR expression in colon cancer cells compared to CLP-SCR (Figure 4H). EGFR inhibitor and CLP-ASO served as positive controls^38^ (Figure 4H and S3H, respectively). To further validate target-specific inhibition, we used EGFR-overexpressing A431 cells, confirming the efficacy of CLP-siEGFR^2’-OMe^ (Figure 4I). Finally, treatment with CLP-siEGFR^2′-OMe^ significantly increased apoptosis in Caco-2 cells, resulting in a reduction in cell survival compared to CLP-SCR (Figure 4J-L).

**Figure 4.**
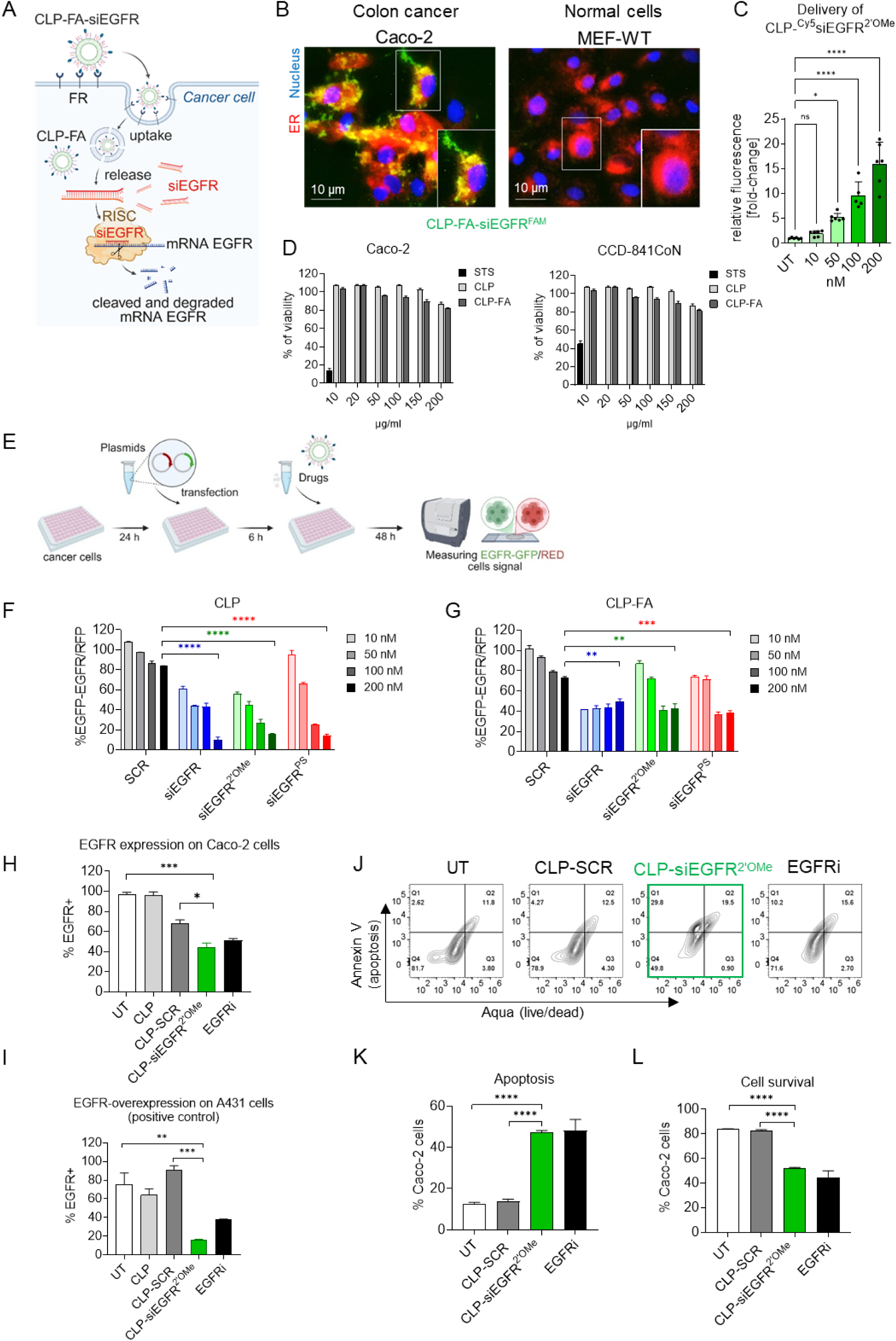
CPL-siRNA complexes achieve targeted EGFR silencing in colorectal cancer cells in vitro. Uptake of CLP-FA-siEGFR via folate receptor (FR)-mediated endocytosis in cancer cells, and the mechanism of action through the RNA-induced silencing complex (RISC) (**A**); Intracellular localization of CLP-FA-siEGFR^FAM^ (100 nM) visualized by fluorescence microscopy (×40 magnification) after 30min incubation with Caco-2 and MEF-WT cells. The nucleus was stained with DAPI (blue), and the endoplasmic reticulum (ER) with ER-Tracker Red (**B**); Delivery of the CLP-^Cy5^siEGFR^2’-OMe^ to assess concentration-dependent uptake (**C**) Cell viability after 48 h incubation with empty CLP and CLP-FA (10–200 µg/mL), compared to the Staurosporine positive control (STS, 1 μM) (**D**); Schematic of the treatment utilizing a DFA tool (pEGFP-EGFR/DsRED) in Caco-2 cells. Cells were co-transfected with pEGFP-EGFR (green) and pDsRED-N1 (red) plasmids using Lipofectamine 2000, then treated with CLP-delivered siRNAs. After 48h, EGFP-EGFR/DsRED fluorescence was measured and normalized to untreated plasmid-transfected controls (C-Control, set to 100%)^35,36,37^ (**E**); Silencing activity of tested EGFR-targeted siRNAs (10-200 nM) delivered via CLP (**F**) and CLP-FA (**G**), in comparison to scrambled (CLP-SCR); Inhibition of EGFR expression by CLP-siEGFR^2′-OMe^ (100 nM) in Caco-2 and A431 cells, assessed by flow cytometry after 48 h incubation (**H-I**); Representative Annexin V/Aqua staining (apoptosis/necrosis) flow cytometry panels of Caco-2 cells treated with CLP-siEGFR^2′-OMe^ (100 nM) for 48 h (**J**), with quantified apoptosis (**K**) and cell survival (**L**). EGFR inhibitor (PD153035, 1 µM) served as a positive control (**H-L**).

### 2.5. CLP-siEGFR^2’-OMe^ is safe and well tolerated in mice

To assess the translation of our CLP-siEGFR^2’-OMe^ in vitro effects into an in vivo setting and to get more insight into the biodistribution and potential off-target toxicity, we monitored the delivery of CLP-siEGFR^2’-OMe^ into the tumors and also different organs. Organ weights of the heart, lungs, liver, kidneys, spleen, and brain showed no significant differences across treatment groups, indicating no apparent organ toxicity (Figure 5A). Serum biochemical analysis revealed stable levels of liver enzymes (CK, ALAT, ASPAT) and kidney function markers (urea, creatinine), suggesting preserved hepatic and renal function (Figure 5B). Additionally, hematological parameters-including red blood cells (RBC), white blood cells (WBC), and platelets (PLT)-remained within normal physiological ranges in PBS group (Figure 5C). These findings confirm that systemic administration of CLP-siEGFR^²′-OMe^ formulation does not induce detectable toxicity, supporting its safety for further preclinical mouse model.

**Figure 5.**
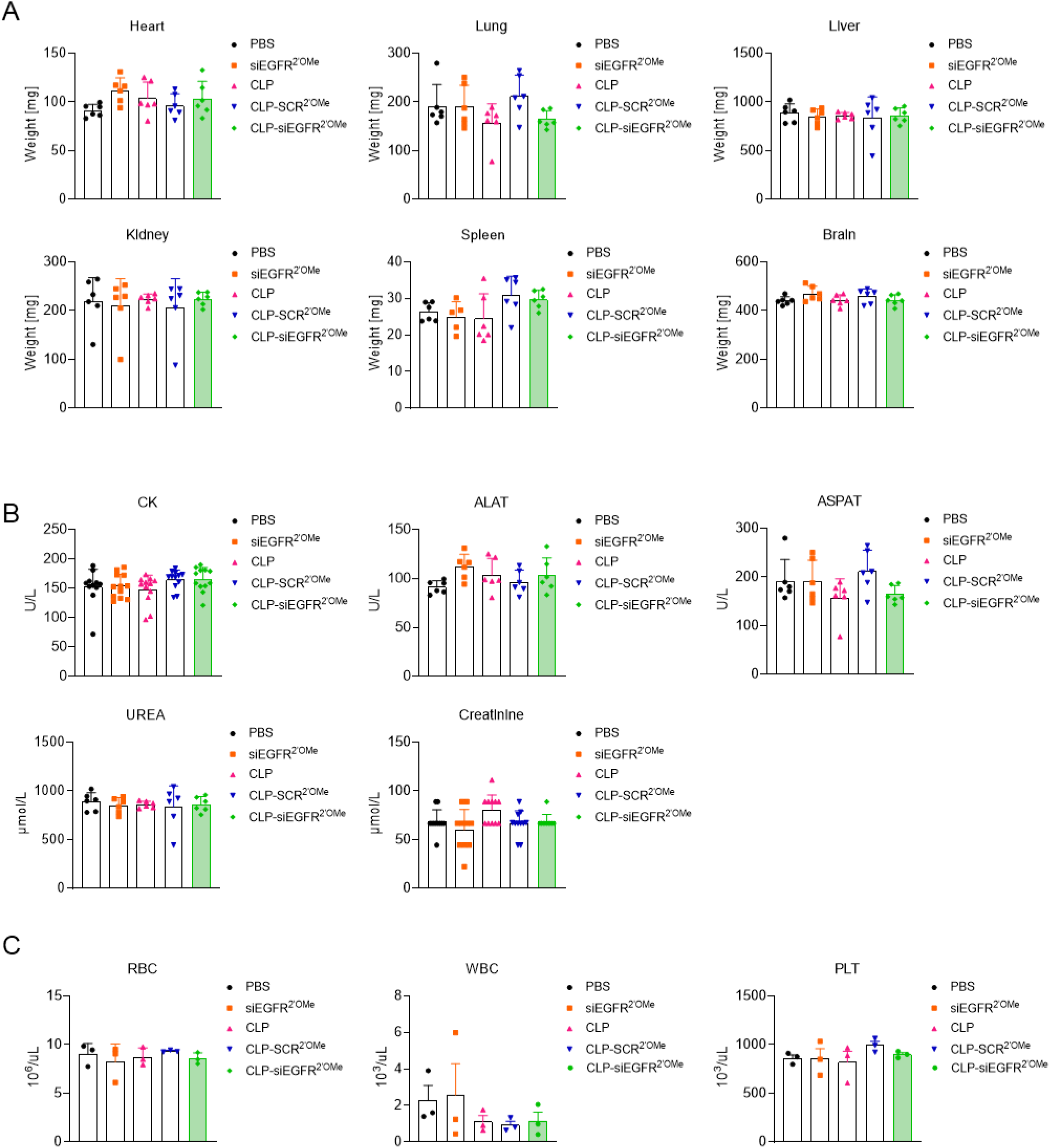
Assessment of biochemical, safety and hemocompatibility after CLP-siEGFR^²′OMe^ treatment in mice. Mice received seven intravenous administrations (i.v.) of the tested compounds at a dose of 2 mg/kg. Analyses were performed two days after the final dose. Organ weights (heart, lung, liver, kidney, spleen, brain) showed no significant changes across treatment groups (**A**); Serum biochemical markers (CK, ALAT, ASPAT, urea, creatinine) remained within normal ranges, indicating no hepatic or renal toxicity (**B**); Hematological parameters (RBC, WBC, PLT) were unaffected, supporting overall safety (**C**). Biochemical parameters (CK: 100-1000 U/L; ALAT: 20-80 U/L; ASPAT: 50-200 U/L; UREA: to 2500-5000 µmol/L; creatinine: 18-53 µmol/L) and hematological parameters (RBC: 7-11 ×10⁶/µL; WBC: 1-4 ×10³/µL; PLT: 500-1200 ×10³/µL) remained within the normal reference ranges for NOG mice. Data shown as mean ± SD (n = 6; female mice). In vivo experiments were conducted in accordance with GLP standards.

### 2.6. CLP-mediated delivery enhances the therapeutic efficacy of siRNA in vivo

Clearance kinetics from the bloodstream significantly influence the rate and extent of tissue distribution, with higher tissue exposure often correlating with increased tissue accumulation.^39^ In this context, we sought to evaluate the impact of the CLP platform on the in vivo clearance kinetics of siRNA, with the objective of enhancing delivery to tumor. A 2 mg/kg dose of either CLP^Cy7^ or CLP-FA^Cy7^ without siRNA^2’-OMe^ cargo, naked ^Cy5^siRNA^2’-OMe^ and CLP-^Cy5^siRNA^2’-OMe^ formulation were intravenously administered to NOG mice bearing Caco-2 xenografts. Whole body imaging was performed at multiple time points post-injection using an IVIS bioluminescence system (Figure 6A). Subsequently, we confirmed that the CLP-^Cy5^siEGFR2^′-OMe^ had enhanced accumulation in tumor xenografts and demonstrated sustained retention over time, as compared to the naked ^Cy5^siEGFR^2′-OMe^, when administered at the same dose (Figure 6B, E, F) and (Figure S4 A, B). Tissue biodistribution studies confirmed efficient delivery of CLP-^Cy5^siEGFR^2′-OMe^ to the intestine with subsequent clearance through the liver and kidneys (Figure S4 C-G).

**Figure 6.**
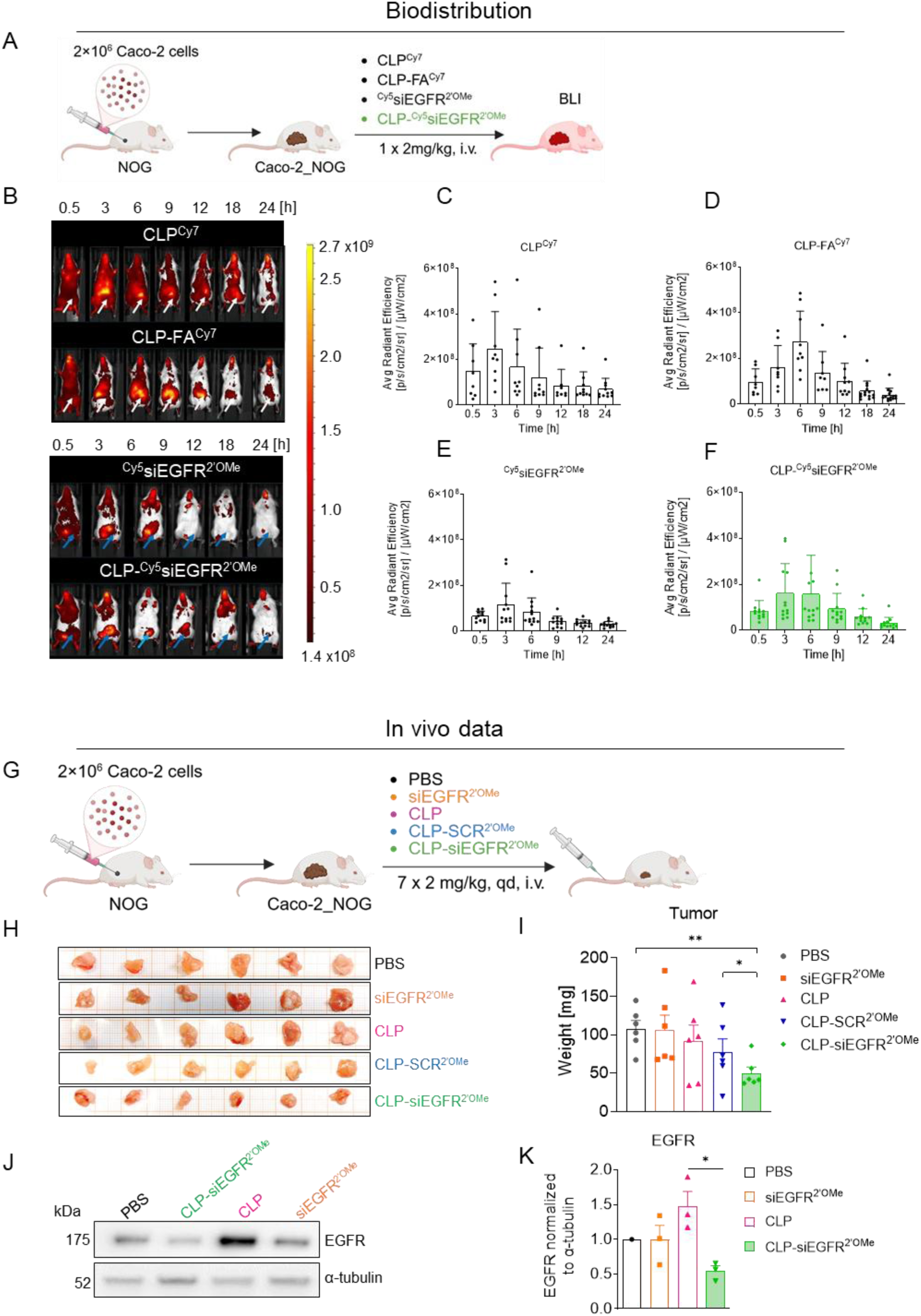
Enhanced in vivo *EGFR* gene inhibition via CLP-enabled siRNA delivery in mice. Graphical study outline illustrating the use of bioluminescence imaging (IVIS) (**A**); Representative IVIS images of mice treated with 2 mg/kg of CLP^Cy7^, CLP-FA^Cy7^, ^Cy5^siEGFR2′-OMe, or CLP^Cy5^siEGFR^2′-OMe^ over time (**B**); Quantitative IVIS analysis shows sustained signal intensity over time following administration of the tested compounds in tumor-bearing NOG mice (**C-F**); Schematic diagram of the treatment regimen: intravenous (i.v.) administration in Caco-2 tumor-bearing NOG mice for 7 consecutive days (**G**); Tumor volume measurements (**H**) and tumor weight at endpoint (**I**); Western blot analysis of EGFR expression and α-tubulin as a loading control (n=3), was performed independently in the Caco-2_NOG model, (**J, K**). Error bars represent mean ± SD (n=6 per group; female mice). In vivo experiments were conducted in accordance with GLP standards.

To determine whether the enhanced tumor accumulation of the CLP-siEGFR^2′-OMe^ formulation translated into functional modulation of target gene expression, mice bearing human colon cancer xenografts were treated intravenously with 2 mg/kg of the tested compounds for 7 consecutive days (Figure 6G). Tumors harvested two days after the final treatment showed a significant mass reduction in the CLP-siEGFR^2′-OMe^ group compared to the scrambled group (CLP-SCR^2′OMe^), as shown in Figure 6H and 6I. Furthermore, western blot analysis confirmed effective target gene silencing, demonstrating a marked decrease in EGFR protein levels in tumors treated with CLP-delivered siEGFR^2′-OMe^ relative to the scrambled control (Figure 6 J, K), but not in the liver (Figure S4 H). In contrast, naked siEGFR^2′-OMe^ demonstrated no measurable impact on EGFR expression or tumor progression. In vivo safety assessment confirmed that new CLP-siEGFR^2′-OMe^ formulation was well tolerated in mice following repeated intravenous administration (2 mg/kg) over 9 days, with no significant changes in body weight (Figure S4I). These findings confirm the efficacy of the CLP platform in mediating targeted siRNA delivery and *EGFR* gene silencing in a preclinical colon cancer model.

## 3. Discussion

The development of effective delivery systems remains the principal barrier to the clinical translation of RNA interference therapeutics for solid tumors.^40^ In this study, we demonstrate that PEGylated cationic liposomes (CLPs), with or without folic acid (FA) functionalization, enable efficient delivery of EGFR-targeted siRNA inhibitor to human colorectal cancer, resulting in robust gene silencing and significant tumor growth inhibition while maintaining a favorable safety profile. Our findings shown that CLP composed of saturated ethylphosphocholine derivatives, neutral phosphatidylcholine derivatives, and PEGylated lipids bearing terminal amine groups can be reproducibly synthesized using thin-film hydration and extrusion (Figure 1 A). Both unmodified and folic acid-modified liposomes (FA-CLPs) exhibited uniform size distribution, low polydispersity, and high stability across a wide range of pH and temperature conditions, during incubation at 37 °C, and under long-term storage (Figure 1 B-F). This physicochemical robustness positions EPC-based systems as a valuable alternative to commonly used DOTAP- and ionizable lipid-based carriers for nucleic acid delivery.^41^

Next, the mechanistic in vitro analyses confirmed efficient siRNA complexation and protection by CLPs, as indicated by gradual ζ-potential reduction, preserved colloidal stability, and validated siRNA binding by PAGE analysis (Figure 3 A-E). Importantly, the complexes remained stable at physiological pH while enabling controlled siRNA release at pH 5.5-6.5 (Figure 3 F, H), consistent with the critical role of endosomal escape in effective RNA delivery.^42,43,24^ Other studies have shown that encapsulation of therapeutic PD-L1 siRNA and an EGFR-targeting short peptide within cationic PEI-LNPs has been reported to enhance stability, enable controlled release, and improve transfection efficiency in EGFR-positive lung cancer.^44^

A key improvement of this work is the incorporation of DSPE-PEG(2000)-NH₂ lipids, which improves colloidal stability and prolongs systemic circulation of the CLP platform. However, PEGylation is associated with the well-known ‘PEG dilemma’, as it can reduce cellular uptake and limit endosomal escape^28,29^. In our system, these effects are mitigated by an optimized lipid composition (Figure 1B), together with ligand-mediated targeting and pH-control release, enabling efficient intracellular delivery despite the presence of a PEG corona. Moreover, this design provided steric stabilization typical of PEGylation while enabling additional electrostatic binding of anionic siRNA without inducing aggregation and while maintaining a net positive surface charge (Figure 3 D, G, E). These findings highlight an underexplored role of PEG end-group chemistry in nucleic acid delivery systems. The observed effects are consistent with evidence showing that functionalized PEG influences surface charge, protein corona formation, and nanoparticle stability in biological environments.^45^ Furthermore, the combined presence of PEG-amine groups and folic acid ligands modulated release kinetics without affecting particle size or disparity, underscoring the importance of surface chemistry in balancing serum stability with pH-dependent release (Figure 3 F, H). Moreover, selective uptake of CLP-FA-siEGFR^FAM^ formulation by Caco-2 human colorectal cancer cells compared to fibroblasts (Figure 4 A, B) validates our targeting strategy based on differential folate receptor (FR) expression. FR is overexpressed in approximately 80% of colorectal adenocarcinomas while showing limited expression in normal tissues, making it an attractive target for selective drug delivery.^46^ Microscopic analysis confirmed cellular uptake within 30 min, consistent with the rapid receptor-mediated endocytosis reported for folate-conjugated nanoparticles (Figure 4 B, S3A).

The concentration-dependent uptake observed with Cy5-labeled CLP-siEGFR^2′-OMe^ (Figure 4 C) and efficient carrier loading (Figure 3 G, S3 B) indicate that chemical modifications do not interfere with CLP formation or cellular internalization. No significant cytotoxicity was observed in cancer cells, normal colon epithelial cells, or macrophages at therapeutic concentrations of empty CLPs (Figure 4 D, S3 C) addresses a longstanding concern regarding cationic lipid formulations in clinical settings.^47^

Incorporation of 2′-*O*-methyl ribose modifications and phosphorothioate internucleotide linkages in siRNA was essential for effective gene silencing in the CLP system (Figure 2 C, D and 4 E-G). Dual fluorescence assays demonstrated that CLP-siEGFR^2′-OMe^ retained approximately 65% target inhibition, comparable to unmodified siRNA (Figure 4 F, G), indicating preserved RISC loading and target recognition.^48^ An in-depth flow cytometry analysis confirmed significant EGFR inhibition in Caco-2 cells (Figure 4 H), likely reflecting altered endosomal trafficking following receptor-mediated uptake. These findings are consistent with reports showing that extensively modified siRNAs can maintain full RNAi activity in vivo.^49^ EGFR-amplified A431 cells were used as a positive control, confirming siRNA specificity and broader applicability of the CLP platform across EGFR-overexpressing cancer models (Figure 4 I). Consistent with previous studies using boron cluster-modified antisense oligonucleotides (B-ASOs) targeting the same EGFR mRNA fragment^38,35,37^, CLP-delivered siEGFR also reduced target expression and promoted apoptosis in cancer cells (Figure 4 F-L). The efficient gene silencing observed with CLP formulations may indicate successful endosomal escape, a key rate-limiting step for lipid-based siRNA delivery (Figure 4 F-I). Although endosomal escape was not directly visualized, functional EGFR inhibition measured by flow cytometry and DFA tool (Figure 4 E-I), confirmed that sufficient siRNA reached the cytoplasm to engage the RNAi machinery. Recently, Mitchell et al. reported the development of an EGFR antibody-conjugated lipid nanoparticle (aEGFR-LNP) platform, engineered to enhance mRNA delivery to EGFR-expressing placental cells for the treatment of pregnancy-related complications.^50^ Agnello et al. investigated the therapeutic potential of anti-EGFR and aptamer-functionalized nanostructures in triple-negative breast cancer (TNBC).^51^ In addition, LNPs were targeted to EGFR (αEGFR-LNP) using a well-established anti-EGFR antibody (cetuximab) to enable delivery of siRNAs against HPV in head and neck squamous cell carcinoma (HNSCC). The resulting αEGFR-LNP-siHPV formulation improved anti-tumor activity, enhanced silencing of HPV targets, and increased apoptosis in EGFR-expressing HNSCC cell lines compared to non-targeted LNPs.^52,24^

Furthermore, biodistribution studies demonstrated favorable pharmacokinetic properties of CLP-siEGFR^2′-OMe^ (Figure 6 A-F). Whole-body imaging revealed clearance over 24 h for both CLP^Cy7^ and CLP-FA^Cy7^, indicating prolonged circulation compared with naked siRNA, which is rapidly eliminated by renal filtration (Figure S4 C-G). Heatmap analysis showed more efficient siRNA release from CLP than CLP-FA at endosomal pH 5.5, likely due to steric hindrance imposed by the folate ligand in vitro study (Figure 3 H). Consistently, CLP-^Cy5^siEGFR^2′-OMe^ exhibited enhanced tumor accumulation in mice and prolonged retention compared with naked siRNA (Figure 6 B-F and S4 A-B), reflecting the combined contributions of passive and active targeting.^53,24^

Significant tumor mass reduction following seven consecutive daily administrations of CLP-siEGFR^2′-OMe^ provides proof-of-concept for therapeutic efficacy (Figure 6 H-I). Western blot analysis confirmed downregulation EGFR protein levels in tumor but not in normal tissue (Figure 6 J-K and S4 H), indicating on-target gene silencing rather than nonspecific effects. The administered dose falls within the range used in clinical siRNA studies, supporting translational relevance.^54^

The absence of body weight loss after treatment (Figure S4 I) and normal organ function markers (Figure 5 A-C) indicate the drug was well-tolerated. This is notable given that hepatotoxicity has limited some lipid nanoparticle formulations with high positive charge density.^55^

Overall, CLP-siEGFR^2′-OMe^ achieves tumor-selective delivery, functional gene silencing, and therapeutic efficacy in a human colorectal cancer xenograft model without detectable systemic toxicity. These findings establish a strong preclinical foundation for lipoplex-mediated siRNA delivery to solid tumors and support further development toward clinical translation.

## 4. Material and Methods

Oligonucleotides were synthesized by phosphoramidite solid-phase method using LCA CPG solid support and commercial phosphoramidites (Glen Research, USA, for phosphoramidite 6-FAM, ChemGenes, USA, for all other phosphoramidites). Hyacinth BMT activator was purchased from empBiotech GmbH (Berlin, Germany), C18 SepPak cartridges from Waters Corp., (Miliford, MA, USA), anhydrous acetonitrile (J.T. Baker brand) and ammonium hydroxide (30%, J.T. Baker brand) from Avantor Performance Materials (Center Valley, PA, USA).

*<u>Negative ion electrospray mass spectra</u>* were obtained on a Synapt G2 Si high-resolution mass spectrometer (Waters) equipped with a quadrupole-time-of-flight mass analyzer (Waters Corp., Miliford,MA, USA). Data collected in a continuous mode was deconvolved using the MaxEnt1 algorithm. *<u>UV absorption measurements</u>* were performed using a Jasco V-770 UV−VIS/NIR spectrophotometer (Jasco Int., Easton, MD, USA). *<u>RP-HPLC analyzes</u>* were performed on a Shimadzu Prominence HPLC system (Kyoto, Japan) using Kinetex 5 µm C-18 column (100 Å, 250 mm × 4.6 mm column, Phenomenex). Analysis conditions for cyanine 5 labelled oligonucleotides s-^Cy5^EGFR^2’-OMe^ and s-^Cy5^scrambled^2’-OMe^: buffer A - 0.1 M CH_3_COONH_4_, pH 6.7; buffer B: CH_3_CN flow: 1 mL / min. Gradient of buffer B: 0 → 2 min 0%, 2 → 25 min 0-48%, 25 → 30 min 48- 60%, 30 → 35 mins 60-0%. UV detection was performed at λmax 260 nm (and at 494 nm for FAM-labelled RNA and at 646 nm for Cy5-labelled RNA), and the amount of compound was defined in optical units (OD) at the wavelength of 260 nm.

*<u>Cationic liposomes</u>*were formulated using 1,2-dimyristoyl-sn-glycero-3-ethylphosphocholine (EPC), 1,2-dipalmitoyl-sn-glycero-3-phosphocholine (DPPC), 1,2-dimyristoyl-sn-glycero-3-phosphocholine (DMPC), and 1,2-distearoyl-sn-glycero-3-phosphoethanolamine-N-[amino(polyethylene glycol)-2000] (DSPE-PEG(2000)-NH₂). For targeted formulations, 1,2-distearoyl-sn-glycero-3-phosphoethanolamine-N-[folate(polyethylene glycol)-2000] (DSPE-PEG(2000)-Folate) was added. Fluorescently labeled liposomes were prepared using 1,2-distearoyl-sn-glycero-3-phosphoethanolamine conjugated with Cyanine 7 (18:0 Cy7 PE). All lipids were purchased from Avanti Polar Lipids (Alabaster, AL, USA) and used without further purification. Ultrapure Milli-Q water (DNase-free and RNase-free grade) was used for all liposome preparation and nucleic acid handling procedures.

### 4.1. Synthesis of RNA oligonucleotides

All RNA oligonucleotides were synthesized at CMMS PAS on 0.1 µmol scale using GeneWorld DNA/RNA synthesizer H6 (K&A Laborgeraete GbR, Schaafheim, Germany) with standard protocols.^56^ The synthesis of oligonucleotides (s-EGFR, as-EGFR, s-scrambled, as-scrambled, s-GFP, as-GFP, s-EGFR^2’-OMe^, as-EGFR^2’-OMe^) was carried out in the DMT-on mode, so that it was possible to separate them as pure 5’-protected RNAs according to the described procedure.^57^ Then, upon removal of the DMT group, oligoribonucleotides were additionally purified by RP-HPLC, desalted on SepPak columns and subjected to the mass spectrometry analysis. For oligonucleotides with phosphorothioate bonds (s-EGFR^PS^, as-EGFR^PS^, s-EGFR^PS/2’-OMe^, as-EGFR^PS/2’-OMe^), the procedure was modified, as the fists step of deprotection/purification, the β-cyanoethyl protecting groups were removed from phosphorus in internucleotide linkages by washing the bed with piperidine in acetonitrile. Subsequent steps followed the standard protocol for the other oligonucleotides. For fluorescently labelled 5’-FAM-antisense strand (as-^FAM^EGFR), 5’-FAM-sense strand (s-^FAM^EGFR) and 5’-Cy5-sense strands (Cy5-EGFR^2’-OMe^, Cy5-scrambles^2’-OMe^) oligonucleotides, the cuts off from the bed were carried out by routine procedure and compounds were directly purified by RP-HPLC (conditions described above). All the retention times and MS data obtained for the synthesized oligonucleotides are summarized in Table 1.

### 4.2. Synthesis of CLP, CLP-FA, CLP^Cy7^, CLP-FA^Cy7^ liposomes

All liposomes were prepared using the thin-film hydration method followed by Bangham et al.,^58^ with our patented components (PCT.PL2022.000046 and WO2023/022615). An appropriate masses of lipids, was dissolved in 2 mL of analytical-grade chloroform. The solution was thoroughly mixed using a vortex mixer. The lipid-chloroform mixture was then transferred to a single round-bottom flask, and the chloroform was evaporated using a rotary evaporator set to 45°C, forming a thin lipid film. Next, 4 mL of sterile saline solution was added to the flask, which was tightly sealed and incubated at 30°C overnight (12 hours) to hydrate the lipid film. After hydration, the lipid layers were vortexed until the entire lipid film detached from the flask walls. The suspension was further incubated for an additional 6 hours at 40°C with continuous shaking. The lipid vesicles were then extruded through a 0.1 µm membrane filter, passing through the membrane 20 times to achieve uniform size. The schematic of the procedure for the preparation of CLPs, CLPs-FA liposomes, and their fluorescently labeled counterparts (CLPs^Cy7^, CLPs-FA^Cy7^) is presented in Figure 1A. The liposome fractions were subsequently combined and dialyzed against ultrapure water for 16 hours using Float-A-Lyzer G2 Dialysis Device membrane MWCO 100kD. The external solution was replaced twice, at 2 and 4 hours. After dialysis, the liposomes were filtered through sterile PTFE syringe filters with a pore size of 0.2 µm. The resulting liposomes were prepared for binding with nucleic acids in subsequent experiments.

### 4.3. Physicochemical characteristic of CLP and CLP-FA liposomes

Dynamic light scattering (DLS) and zeta potential analyses were performed using a Zetasizer Ultra instrument (Malvern Panalytical, UK) equipped with ZS Xplorer software. Measurements were conducted at 25 °C after 30 s equilibration. For size and polydispersity analysis, samples were placed in disposable sizing cuvettes (ZEN0040), with the material type set to “liposomes” and dispersant set to water. Data were processed using the *multiple narrow modes* algorithm. For zeta potential measurements, 800 µL of liposome suspension was transferred into a folded capillary zeta cell (DTS1070, Malvern Panalytical). Measurements were carried out under identical conditions (25 °C, 30 s equilibration).

### 4.4. Preparation of CLP-siRNAs for measurements

Cationic liposomes (CLP and CLP-FA) were prepared as described above and dialyzed against ultrapure water. Unmodified siRNA (siEGFR) and chemically modified siRNA (siEGFR^2’-OMe^) were mixed with the liposomes to obtain defined siRNA:CLP mass ratios of 1:100, 1:50, 1:20, and 1:10 (w/w). The mixtures were incubated for 30 minutes at room temperature to allow complex formation. Unless otherwise noted, measurements were performed at 25°C. Two formulation variants were analyzed in parallel (CLP and CLP-FA).

### 4.5. Formation of [^32^P]-siRNA-lipoplex complex analyzed by PAGE

To 0.1 OD of the RNA strands (sense and antisense) dissolved in 15 µL of Milli-Q water [^32^P-γ] ATP (37.0 MBq, 1.00 mCi), T4 polynucleotide kinase (1 µL, 10,000 units / mL), and 2 µL of phosphorylation reaction buffer supplied by the kinase manufacturer (final volume 20 µL). The reaction mixture was then incubated at 37°C for 1 hour. The next step was to inactivate the kinase by incubating the reaction mixture at 85°C for 3 minutes. [^32^P]-siRNA was assembled by combining solutions of each strand of [^32^P]-RNA radiolabelled in a 1: 1 molar ratio in sterile milliQ water to a final volume of 20 µL. The mixture was heated at 85°C for 5 min, then slowly cooled to room temperature for 1 h. The radiolabeled duplex [^32^P]-siRNA solution was added to the lipoplex solution (CLP, CLP-FA) in the defined weight ratio (1:1, 1:2.5, 1:10, 1:20), followed by incubation at room temperature for 30 min. After this incubation time, the samples were applied to a 15% polyacrylamide gel (PAGE) under non-denaturing conditions (without the addition of urea). The electrophoresis was performed at room temperature with a voltage of 300-400 V / cm and a current of 60 mA for 30 minutes. Then, electrophoresis gels were developed autoradiographically on diagnostic membranes.

### 4.6. Preparation of pH buffers for [^32^P]-siRNA release studies from lipoplex complexes

A buffer with pH 4.0 was prepared based on NaOAc sodium acetate and acetic acid (150 mM) in 10 ml mQ water. NaOAc was combined with acetic acid to obtain a buffer pH 4.0. In this case, it was not possible to use the buffer prepared on the basis of PBS, due to the different buffer capacity of this solution. The remaining buffers (5.5, 6.0, 7.4) were made based on Na_2_HPO_4_ and NaH_2_PO_4_ (150 mM) in 10 ml mQ water. By combining the two solutions in different proportions, buffers with a higher pH were obtained (5.5, 6.0, 7.4). The pH of the solutions was confirmed by the potentiometric method by measuring with a pH-meter.

### 4.7. Release of [^32^P]-siRNA cargo from the formulation

Labeled ^32^P-siRNA was coated with lipoplex (CLP, CLP-FA) in a mass ratio of 1:10, incubation 30 minutes, at room temperature. Then added to the [^32^P]-siRNA (controls) alone as well as the coated systems. In the next step, the samples were incubated with pH buffer (4.0, 5.5, 6.0, 7.4) for 30 minutes, 1 hour, 2 hours and 4 hours at room temperature. Then, mixed with loading buffer (without EDTA) was added to the samples and a double PAGE analysis was performed from the anode (+) to the cathode (-) and inversely as well.

### 4.8. Stability studies of siRNA against lipoplex after 7-day incubation

Unmodified siEGFR and scrambled labeled and assembled according to the above procedures were coated with CLP lipoplex in the ratio 1:10 followed by incubation for 30 minutes at room temperature. After this time, the samples were incubated at 37°C or 4°C, and after 7 days their electrophoretic analysis was performed (15% PAGE, non-denaturing conditions) either in the form of a siRNA-lipoplex complex or after release of oligonucleotides (from the siRNA-lipoplex formulation after the addition of 10-fold excess, i.e. 15 µL of phosphate buffer at pH 5.5). Prior to loading on the gel, 5 µL of loading buffer without EDTA was added to the samples and electrophoretic analysis was performed.

### 4.9. Cell viability

Caco-2 and CCD-841CoN cells were seeded in 96-well plates and incubated overnight to allow attachment. The culture medium was then replaced with fresh medium containing varying concentrations of empty CLP or CLP-FA lipoplex (ranging from 10 to 200 µg/mL). Staurosporine (1 µM) was used as a positive control for cell death. After 48 h of incubation, the MTT assay was performed as described in our previous work.^35^ Viability of differentiated THP-1 and RAW 264.7 macrophages was assessed using MTS assay. Cells were treated with lipoplexes as described above. Cell viability was determined using the CellTiter 96 AQueous One Solution Cell Proliferation Assay (MTS) from Promega (Madison, WI, USA). Following 24 hours of treatment, 10 µL of the MTS reagent was added directly to each well containing 100 µL of culture medium. The plates were incubated at 37°C for 3 h in a humidified, 5% CO2 atmosphere. The absorbance of the soluble formazan product was measured at 490 nm, with a read at λ = 640 for background subtraction, using a BioTek Cytation 5 platform (Agilent, Santa Clara, CA, USA). For both assays, cell viability was calculated as a percentage relative to the untreated control cells (set to 100%).

### 4.10. Assessment of inhibitory activity of siRNAs coated with CLP lipoplexes

CaCo-2 and A431 cells were cultured according to the ATCC procedure. On the day before microscopic observations, cells were seeded in black 96-well plates (clear bottom) at 15 × 10^3^ cells in 100 μL of complete medium per well and left overnight at 37°C and 5% CO_2_. After overnight incubation, full medium was removed from the cells and replaced with OPTI-MEM base medium. Transfection was carried out using commercial Lipofectamine 2000 reagent (Invitrogen) in a ratio of 2:1 (2 µL Lipofectamine 2000 per 1 µg nucleic acid) according to manufacturer’s protocol. In the dual fluorescence assay (DFA) CaCo-2 cells were transfected with pDsRed-N1 reporter plasmid (30 ng/well) and pEGFP-EGFR plasmid (100 ng/well). Then, appropriate siRNAs (10-200 nM) were co-transfected, delivered to cells with the tested lipoplexes CLP or CLP-FA or as a control with a commercial transfection reagent (Lipofectamine 2000) according to the manufacturer’s protocol. After 5 hours of incubation, the transfection mixture was withdrawn from each well and cells were flooded with 200 µL of fresh culture medium containing antibiotics. After 48 h of incubation at 37 °C in 5% CO_2_, cells were washed twice with PBS buffer (no Ca^2+^and Mg^2+^) and lysed overnight with NP-40 buffer (150 mM NaCl, 1% IGEPAL, 50 mM Tris-HCl pH 7.0 and 1 mM PMSF) at 37°C. Prepared cell lysates were used to determine the level of fluorescence were determined using a FLUOstar Omega reader (BMG LABTECH). Flow cytometry was performed as previously described.^38^

### 4.11. Western Blotting

Total protein was extracted from excised equivolume tissue samples. Tissues were homogenized in ice-cold 1 × Cell Lysis Buffer [Cell Signaling Technology (CST), Danvers, MA, USA] supplemented with a phenylmethanesulfonyl fluoride (1 mM, Merck) and 1 × Protease and Phosphatase Inhibitor Cocktail [Thermo Fisher Scientific (TFS), Waltham, MA, USA] using a using Bio-Gen PRO200 Homogenizer (PRO Scientific, Oxford, CT, USA). The lysates were centrifuged at 18,188 × g for 10 min at 4°C. Protein concentrations were determined in the obtained supernatants using the BCA Protein Assay (TFS). Equal amounts of protein were denatured at 95°C for 5 min in Laemmli sample buffer, separated by 8-12% PAGE in MES-SDS buffer (90 V for 10 min and 120 V for 60 min), and transferred onto PVDF membranes (TFS) using the iBlot2 (TFS) system (20 V for 1 min, 23 V for 4 min, and 25 V for 2 min). Membranes were blocked with 3% non-fat dry milk (Merck) in Tris-buffered saline with 0.1% Tween 20 (TBST) for 1 h. After a single rinse with 1 × TBST, the membranes were incubated overnight at 4°C with primary antibodies against EGFR (rabbit, AbCam, Cambridge, UK, #AB52894) or α-tubulin (rat, TFS, #MA180017). Following four washes with TBST, membranes were incubated with anti-rabbit [HRP-conjugated from CST, #7074] or anti-rat (AlexaFluor555-conjugated, #4417) secondary antibodies. After 1 h of incubation at room temperature the membranes were rinsed with 1 × TBST. All antibodies were diluted in 1 × TBST with 3% BSA at 1:2000 for primaries and at 1:10000 for secondaries.

When necessary, protein bands were visualized using Westar Supernova ECL reagent (Cyanagen, Bologna, Italy). Image acquisition was performed on an Azure c400 (Azure Biosystems Inc., Dublin, CA, USA). Densitometric analysis was performed on sub-saturated images using ImageJ 1.54f software (National Institutes of Health, Bethesda, MD, USA), with EGFR band intensities normalized to the loading control (α-tubulin).

### 4.12. In vivo study

Six-week-old female NOG-F mice (strain NOD.Cg-Prkdc^scid Il2rg^tm1Sug/JicTac) were obtained from Taconic Biosciences and used for xenograft experiments. This strain represents a highly immunodeficient model lacking functional T, B, and NK cells and is widely used for human tumor xenograft studies. Upon arrival, animals were subjected to a 7-day quarantine, followed by handling and habituation, in full compliance with GLP standards prior to the initiation of experimental procedures.

Mice were housed in groups of five per cage under specific pathogen-free conditions with controlled environmental parameters (temperature 20–24 °C, relative humidity 45–65%, 12 h light/dark cycle, and 15 air changes per hour). Animals had free access to a standard rodent diet (Altromin) and water ad libitum. Environmental enrichment, including nesting material and shelters, was provided to promote natural behaviors and reduce stress. All animal procedures were performed in accordance with national regulations for the care and use of laboratory animals and were approved by the Local Ethical Committee for Animal Experiments (approval no. 90/2023). For xenograft implantation, Caco-2 cells were detached at logarithmic growth phase using trypsinization, collected and centrifuged to obtain a pellet. The cell pellet was resuspended in sterile phosphate-buffered saline (PBS). Cell number and viability were determined using a hemocytometer and the suspension was adjusted to the required concentration. Immediately prior to implantation, cells were prepared at a density of 2 × 10⁶ cells per injection in PBS and kept on ice until administration.

Female NOG mice bearing Caco-2 xenografts were monitored until tumor volumes reached 50-100 mm³. Mice were randomly assigned to five groups (n = 6 per group): PBS, naked siEGFR^2’OMe^, empty CLP, CLP-SCR^2’OMe^ and CLP-siEGFR^2’OMe^. Treatments were administered via tail vein injection of siRNA dose of 2 mg/kg body weight and ten-fold more for CLP. Groups receiving treatment were administered formulations three times at defined intervals of 48h. Body weight and tumor volume were recorded at each administration and at subsequent time points. Forty-eight hours after the last injection, animals were euthanized by decapitation. Blood (200 µL) was collected into EDTA-containing tubes for immediate hematological analysis. Samples were centrifuged (3000 rpm, 10 min, 4 °C) to separate plasma; 50 µL plasma was frozen at -80 °C, while the remainder was used for biochemical analyses. Organs, including brain, heart, lungs, liver, kidneys, and spleen, were excised, weighed, and frozen on dry ice. Tumors were excised, weighed, photographed, and frozen for subsequent analyses.

### 4.13. Treatment and in vivo imaging

Female NOG mice bearing Caco-2 xenografts were randomly assigned to four treatment groups (n = 8): CLP^Cy7^, CLP-FA^Cy7^, ^Cy5^siRNA^2’-OMe^ and CLP^Cy5^siRNA^2’-OMe^. The dosing was 2 mg/kg body weight for siRNA and ten-fold higher for CLP formulations. Control animals were left untreated and were included solely as reference to assess background tissue auto-fluorescence. This allowed accurate interpretation of fluorescence signals from experimental groups. Formulations were administered via tail vein injection. In vivo fluorescence imaging was performed using an IVIS Spectrum (Perkinelmer) imaging system at wavelengths appropriate for Cy5 and Cy7 dye. Tumor and biodistribution imaging were conducted at 0, 6, 12, 18 and 24 hours post-injection. Mice were anesthetized with isoflurane (3–4% induction, 1.5% maintenance, oxygen flow 1 L/min) during imaging. Images were acquired from both dorsal and ventral sides, and tails were covered to eliminate fluorescence from the injection site.

### 4.14. Blood Hematology and Serum Biochemistry Assessment

Blood samples were collected from euthanized mice into EDTA-containing tubes. Hematological analysis was performed using an automated hematology analyzer Abacus Junior Vet 5 following the manufacturer’s instructions. The following parameters were evaluated: White Blood Cells (WBC), Red Blood Cells (RBC) and Platelets (PLC). All measurements were performed in duplicate, and instrument calibration and quality controls were verified prior to analysis to ensure accuracy and reproducibility.

Plasma was separated by centrifugation at 3000 rpm for 10 minutes at 4 °C, and serum was collected for biochemical analysis. Serum markers were quantified using commercial kits from Cormay following the manufacturer’s protocols. All measurements were performed on a UV–Vis plate reader (Synergy H1, BioTek) with appropriate blanks and calibration curves.

Creatine Kinase MB (CK-MB): CK-MB activity was determined using the Liquick Cor-CK-MB 30 kit (Cormay). The assay is based on a coupled enzymatic reaction in which CK-MB catalyzes the conversion of creatine phosphate and ADP to creatine and ATP. The generated ATP is subsequently utilized in auxiliary reactions leading to NADPH formation, and the change in absorbance at 340 nm is measured spectrophotometrically.

Alanine Aminotransferase (ALAT): ALAT activity was measured using the Liquick Cor-ALAT 60 kit (Cormay). The method is based on the conversion of alanine and α-ketoglutarate to pyruvate and glutamate. Pyruvate is subsequently reduced to lactate in the presence of lactate dehydrogenase, accompanied by oxidation of NADH to NAD⁺. The decrease in absorbance at 340 nm is proportional to ALAT activity.

Aspartate Aminotransferase (ASAT): ASAT activity was determined using the Liquick Cor-ASAT 60 kit (Cormay). The reaction involves transamination of aspartate and α-ketoglutarate to oxaloacetate and glutamate. Oxaloacetate is subsequently reduced in a coupled enzymatic reaction involving NADH, and the decrease in absorbance at 340 nm is monitored.

Urea: Urea concentration was measured using the Liquick Cor-UREA 60 kit (Cormay). In this enzymatic assay, urea is hydrolyzed by urease to ammonia and carbon dioxide. The produced ammonia participates in a subsequent enzymatic reaction generating a chromogenic product, which is measured spectrophotometrically at 520–550 nm.

Creatinine: Creatinine levels were determined using the Liquick Cor-CREATININE 60 kit (Cormay). The assay is based on the Jaffe reaction, in which creatinine reacts with picrate ions in an alkaline environment to form a colored complex. The intensity of the color, measured spectrophotometrically at 490–510 nm, is proportional to the creatinine concentration.

All assays were calibrated and validated prior to analysis using appropriate calibration and quality control reagents, including CORMAY CK-MB CALIBRATOR, CORMAY MULTICALIBRATOR LEVEL 1, LEVEL 2 and quality control sera (CORMAY SERUM HP, CORMAY SERUM HN). These reagents were used to verify the accuracy and reliability of the analytical methods according to the manufacturer’s recommendations. All measurements were performed in duplicate to ensure reproducibility.

### 4.15. Statistical Analysis

An unpaired Student t-test was used to calculate 2-tailed P-values to estimate statistical significance between 2 treatment groups. One-way analysis of variance and Bonferroni post-test were used to assess differences between multiple groups and in tumor growth kinetics, respectively. Statistically significant *P*-values were indicated in the figures compared to untreated or PBS groups). Data are presented as mean±SD (n=3-6); *P < 0.05, **P < 0.01, ***P < 0.001, ****P < 0.0001 by one-way ANOVA with Bonferroni’s correction post hoc test. Data were analyzed using Prism software version 10 (GraphPad Software).

## 6. Acknowledgments

This research was funded by the Medical Research Agency (Poland) under the project implemented by Biotechna S.A., grant no. 2022/ABM/06/00005. The authors thank Anna Maciaszek (CMMS PAS) for the synthesis of oligonucleotides, Ewelina Wielgus (CMMS PAS) for the mass spectrometric analysis.

## Supplementary

**Figure S1.**
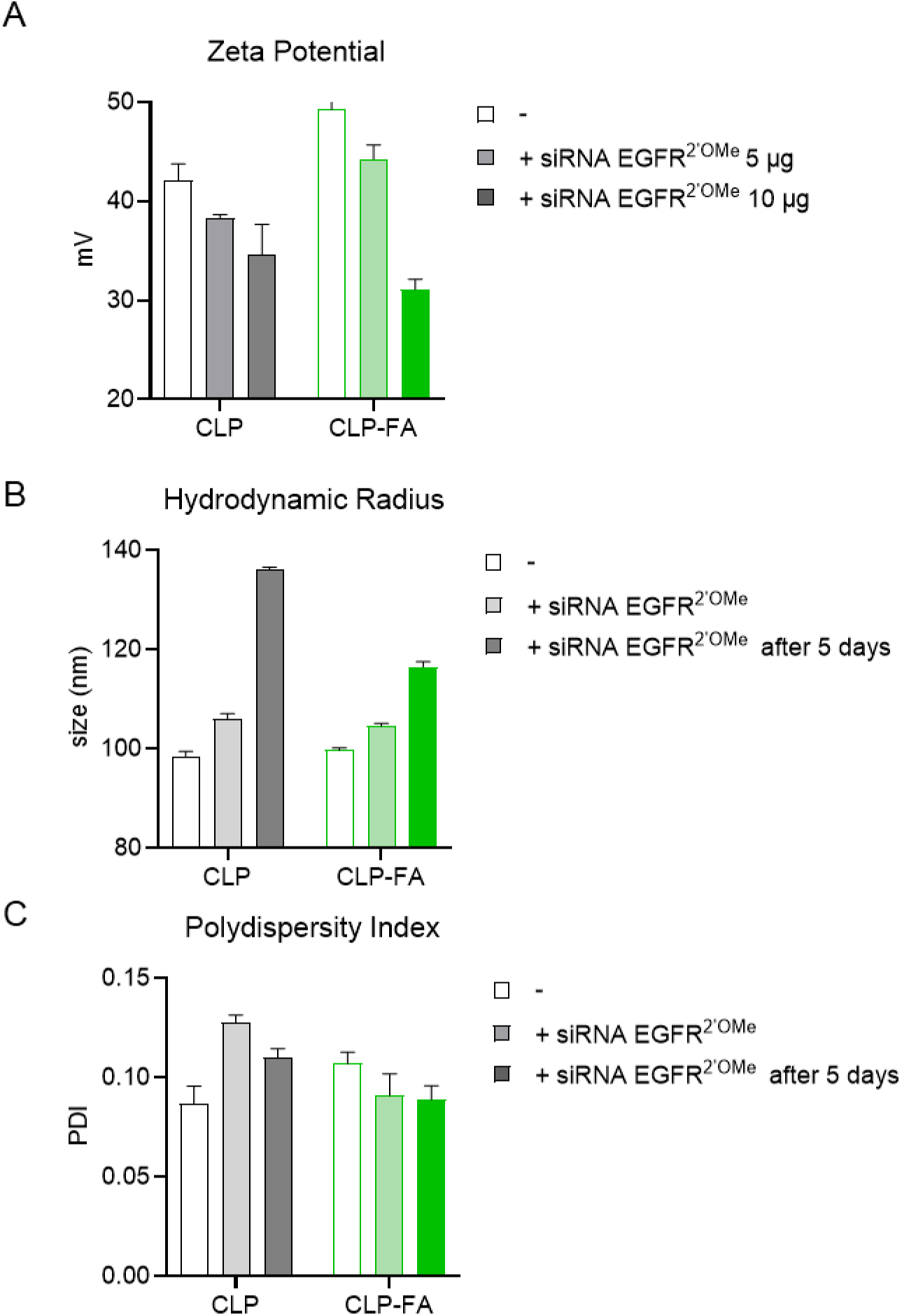
Physicochemical characterization of CLP and CLP-FA lipoplexes with and without siRNA. Zeta potential (**A**); hydrodynamic radius (**B**); and polydispersity index (PDI) (**C**) of empty lipoplexes and lipoplexes loaded with siRNA EGFR^2′OMe^ (5 or 10 μg, as shown), measured immediately after preparation and after 5 days of storage.

**Figure S2.**
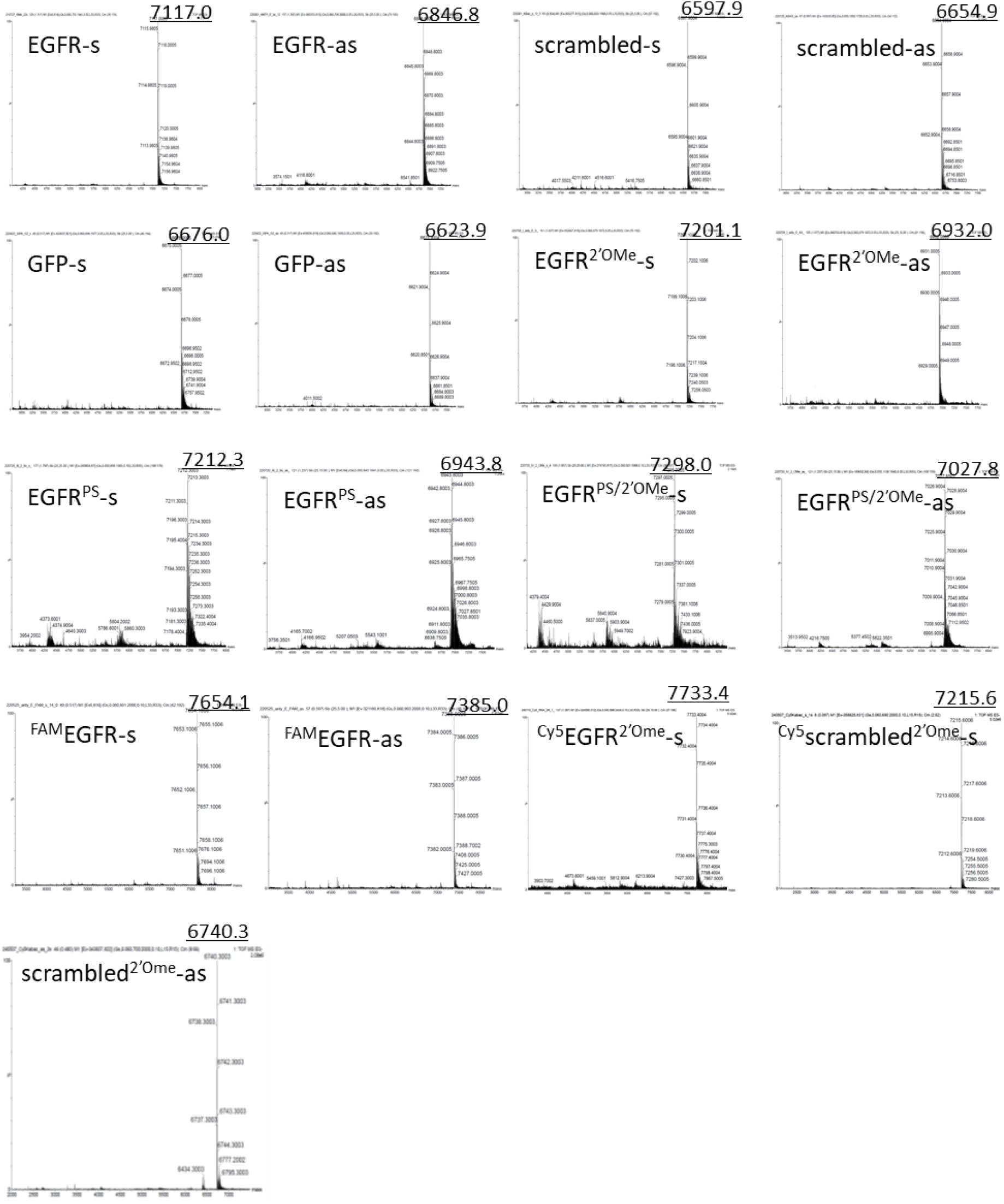
ESI-Q-TOF mass spectra of unmodified and chemically modified siRNA strands (sense and antisense), including PS, 2′-*O*-methyl, and PS/2′-O-methyl variants, as well as fluorophore-labeled oligonucleotides. Measured masses are consistent with theoretical values.

**Figure S3.**
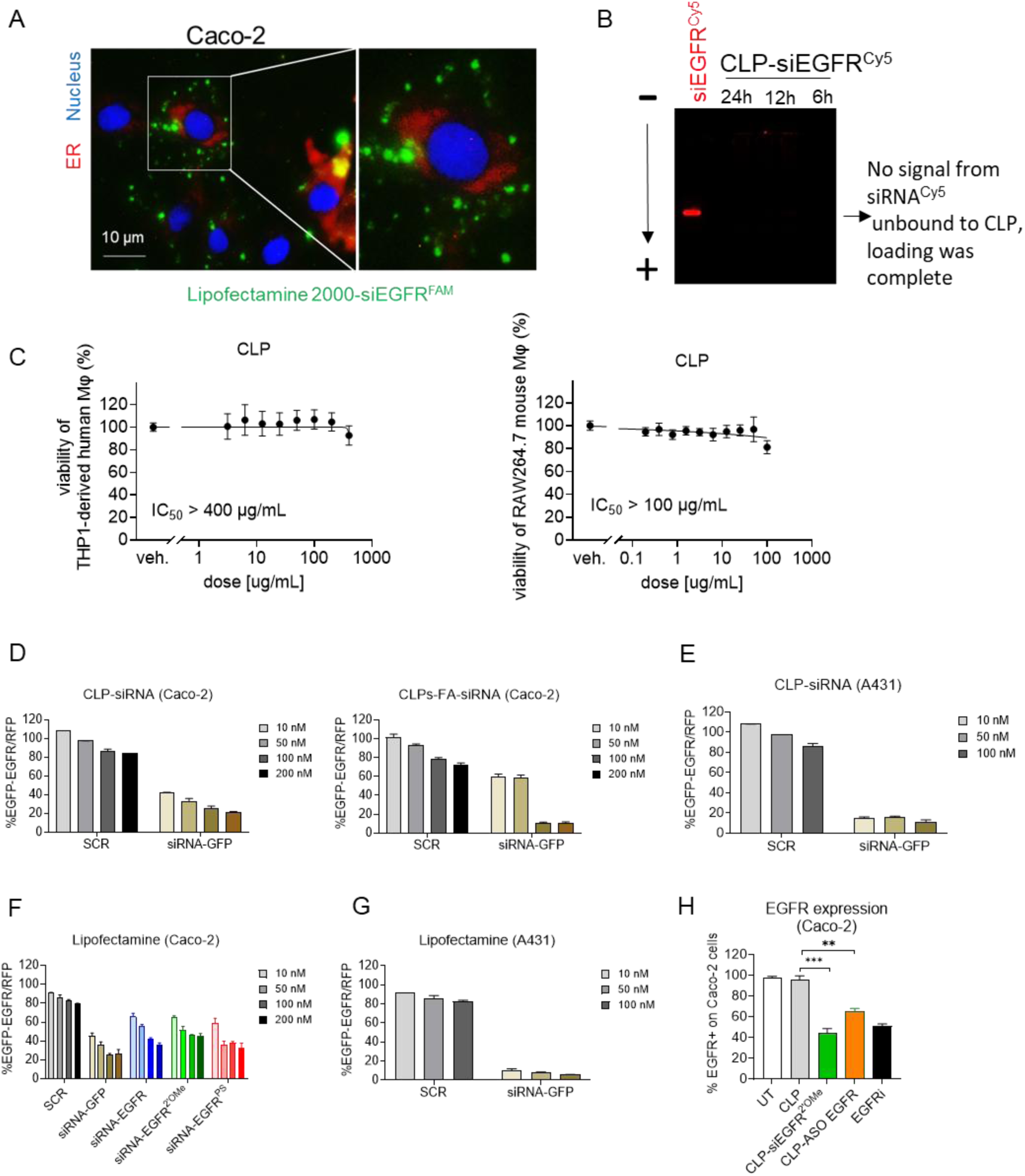
Properties and Activity of EGFR-targeting siRNA lipoplexes. Fluorescence microscopy images (×40 magnification) of Caco-2 cells treated with Lipofectamine 2000-complexed siEGFR^FAM^ (100nM, green dots), demonstrating intracellular localization after 30min incubation; nuclei stained with DAPI (blue) and the endoplasmic reticulum stained with ER-Tracker RED (**A**); native 15% PAGE analysis confirming complete loading of Cy5-labeled siEGFR into CLP lipoplexes (**B**); cell viability of THP-1-derived human macrophages and RAW264.7 mouse macrophages following CLP (0.1-200 µg/mL) treatment with IC50 value, assessed after 48 h by MTT assay (**C**); control GFP silencing in Caco-2 and A431 cells treated with CLP- or CLP-FA-formulated siRNA-GFP at the indicated concentrations by DFA (**D-E**); EGFR silencing mediated by Lipofectamine 2000-formulated siRNAs in Caco-2 and A431 cells after 48 h by DFA tool (**F-G**); EGFR expression on Caco-2 cells following treatment with empty CLP, CLP-siRNA EGFR, and CLP-ASO EGFR formulations (100 nM, 48 h; positive controls), compared with the EGFR inhibitor PD153035 (1µM), as determined by flow cytometry.

**Figure S4.**
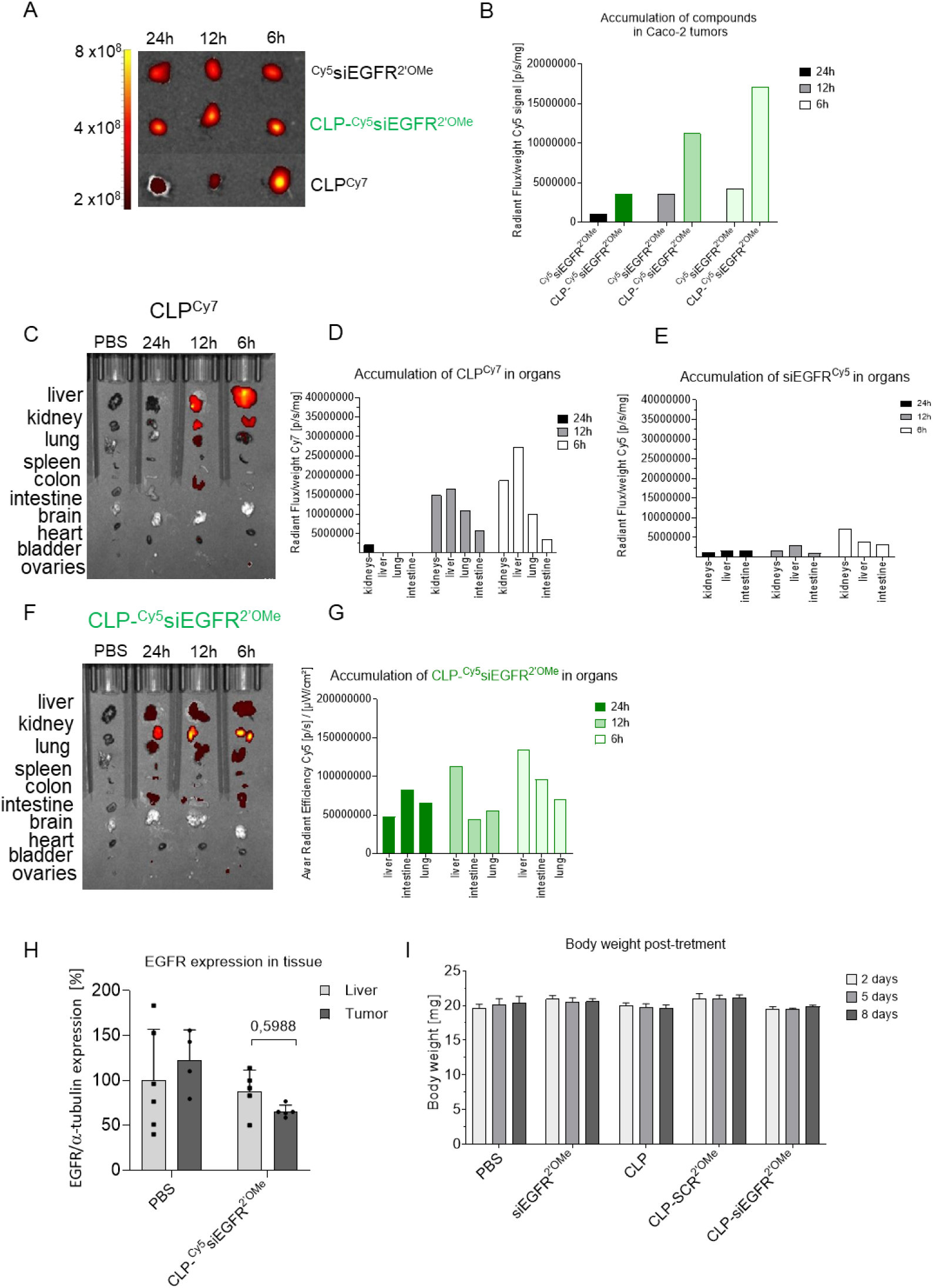
In vivo biodistribution and tumor accumulation of CLP-siEGFR formulations. Quantification of siRNA ^Cy5^EGFR^2’-OMe^, ^Cy5^EGFR^2’-OMe^ encapsulated in CLP, and CLP alone in tumor tissue at 6, 12, and 24 hours post-intravenous injection (2mg/kg) by IVIS (**A**); Comparative accumulation of CLP-^Cy5^EGFR^2’-OMe^ and siRNA ^Cy5^EGFR^2’-OMe^ (2mg/kg) in tumors at the indicated time points (**B**); Biodistribution of empty CLP^Cy7^ in major organs of mice at 6, 12, and 24 hours post-injection (**C-D**); Organ-specific accumulation of siEGFR^Cy5^ at the same time points (**E**); Biodistribution and accumulation of CLP-^Cy5^EGFR^2’-OMe^ (2mg/kg) in mouse organs over 6, 12, and 24 hours by IVIS (**F-G**); EGFR expression in liver and CaCo-2 xenografts, assessed by western blot, following treatment with indicated compounds (2 mg/kg, intravenous) (**H**); Body weight of mice monitored post-treatment, indicating systemic tolerability (**I**). Data are presented as mean ± SD.

